# Nursery habitat use by juvenile blue crabs in created and natural marshes

**DOI:** 10.1101/2020.07.10.197830

**Authors:** D.M. Bilkovic, R.E. Isdell, D. Stanhope, K.T. Angstadt, K.J. Havens, R.M. Chambers

## Abstract

1. Climate change and coastal development pressures have intensified the need for shoreline protection. Nature-first approaches that use natural habitats, particularly marshes, are being promoted globally as ecologically-beneficial alternatives to grey infrastructure. The ability of these novel shorelines to provide nursery habitat to blue crab, an ecologically and economically important species along the Atlantic and Gulf coasts of the United States, has not been quantified.
2. We quantified the abundance and size distribution of juvenile blue crabs from a chronosequence of living shorelines (created fringing marshes) spanning 2 to 16 years in age (since construction) and compared with paired natural fringing marshes in the southern Chesapeake Bay.
3. Both created and natural fringing marshes are being used by blue crabs as primary nursery habitats. While there were interannual differences in abundance, young blue crabs (≤ 2.5 cm carapace width) were observed in similar densities and sizes at living shoreline and natural marshes. There was no relationship between the age of the living shoreline and blue crab density, indicating that even the youngest living shorelines (2 years) were providing primary nursery habitat. Young blue crabs were more abundant in more isolated marshes and those that were inundated for longer periods of time each tidal cycle, which may be evidence for habitat-limitation.
4. *Synthesis and applications:* We provide evidence that juvenile blue crabs are comparably using natural and created fringing salt marshes as primary nursery habitat. Although the relative importance of salt marshes as young crab nursery habitat is not fully understood and likely varies by system, the value of marshes within a suite of available structural nursery habitats may increase under a changing climate. The potential for living shorelines to serve as nursery habitat for an economically important species may provide additional incentives to implement these climate adaptation strategies.

## INTRODUCTION

Blue crab (*Callinectes sapidus*) have a complex life history that includes a series of ontogenetic shifts of estuarine or coastal habitat use during their juvenile stages. Larvae develop in open water offshore environments and megalopae (postlarvae) enter inshore estuarine nurseries using tidal currents (Forward, Tankersley, & Welch, 2003; Epifanio 2019). Postlarvae then settle into a structural primary nursery habitat, such as seagrass beds or salt marshes, that provide refuge and forage opportunities during this vulnerable early growth period (Heck, Hays, & Orth, 2003; Minello, Able, Weinstein, & Hays, 2003; Epifanio 2019). Larger juveniles (>2.5 cm carapace width (cw)) may then disperse throughout the estuary to multiple secondary nursery habitats, including unstructured muddy habitats adjacent to salt marshes (King, Hines, Craige, & Grap, 2005; Seitz, Lipcius, & Seebo, 2005). The importance of different primary nursery habitats for postlarvae and young juvenile blue crabs (≤2.5 cm cw) is not fully understood and may vary by region, with some previous studies describing seagrass as being preferred in Chesapeake Bay (Heck and Thoman, 1984; Orth and Van Montfrans, 1987) and salt marshes being preferred in estuaries of Delaware, North Carolina and the Gulf of Mexico, among others (Etherington & Eggleston, 2000; Jivoff & Able 2003; Minello et al., 2003; Minello & Webb, 1997). However, historically, declines in seagrass habitat were not reflected in declines in blue crab populations in Chesapeake Bay (Hines, 2003), and more recent recovery of seagrass beds (since 2012, Lefcheck, Wilcox, Murphy, Marion, & Orth, 2017) has not been matched by concomitant increases in blue crab populations (CBSAC 2019), which suggests additional nursery habitats are being used by young crabs. In addition, most studies evaluated the importance of seagrass for juvenile fishes and invertebrates, including decapod crustaceans, relative to bare unstructured habitats (Minello, Able, Weinstein, & Hays, 2003; McDevitt-Irwin, Iacarella, & Baum, 2016) and the number of studies on seagrass nursery habitat far exceed those on other structural habitats (Lefcheck et al., 2019). Further complicating habitat comparisons is that salt marshes vary in type (fringing, extensive), tidal amplitude, and are composed of multiple habitats (e.g., vegetated edge, inner marsh, intertidal creeks) which can affect juvenile habitat use (Minello, Able, Weinstein, & Hays, 2003).

Young juvenile blue crab (≤ 2.5 cm cw) use of salt marsh habitats, particularly fringing shoreline marshes, as primary nurseries is not well-enumerated in Chesapeake Bay. Maintenance of these habitats is threatened by the interacting pressures of sea level rise and human development (Mitchell, Herman, Bilkovic, & Hershner, 2017). In addition, new shoreline management approaches that use created fringing marsh as a form of protection (living shorelines henceforth) are being encouraged and implemented throughout the geographic range of blue crabs. Living shorelines are preferred because, unlike traditional armoring (bulkhead, riprap), they maintain coastal processes, restore shoreline habitats, and have greater potential to be resilient to sea level rise (Gittman, Popowich, Bruno, & Peterson, 2014; Sutton-Grier, Wowk, & Bamford, 2015; Bilkovic et al. 2017; Smith et al., 2017; Mitchell & Bilkovic, 2019). There is little to no information on the use of these created marshes by young blue crabs as nursery habitat. An additional unknown is the extent to which the use of both natural and created marsh habitat may be mediated by the shorescape setting (shoreline zone that includes riparian, intertidal and the littoral areas of the nearshore waters), including the amount of connected wetland habitat and shoreline armoring. Understanding the relative importance of natural and created fringing salt marshes as young blue crab nursery habitat, in different shorescapes, could be used to inform species management, restoration targeting, and shoreline management.

Here, we (1) comparatively evaluate the abundance of juvenile blue crabs in paired created and natural fringing salt marshes within different landscapes, and with different degrees of connectivity with surrounding shoreline marshes in Chesapeake Bay, and (2) estimate the average time (years) necessary for habitat use of living shoreline marshes by juvenile blue crabs to match the observed use of natural fringing marshes used as reference wetlands. We hypothesize that young blue crabs comparably use created and natural fringing marshes as nursery habitats and that young blue crab abundance increases with the degree of connectivity of the shorescape setting. In addition, we hypothesize that the time to equivalent use of created marshes and reference marshes by these mobile fauna is relatively short (<3 years since marsh construction).

## MATERIALS AND METHODS

Juvenile blue crab density was measured from a chronosequence of living shorelines (created fringing marshes) spanning 2 to 16 years in age (since construction) and compared with paired natural fringing marshes in the southern Chesapeake Bay (**Figure 1**). From an initial candidate pool of more than 100 living shorelines with similar design features (created marsh, sand fill, and stone sill) that was extracted from the Virginia Shoreline Permit Database (CCRM 2018), 13 sites were selected on the basis of 1) presence and accessibility of a paired natural fringing marsh within close proximity and with similar environmental conditions (e.g., exposure, land use, salinity), 2) property-owner permissions, 3) age of project (minimum of 2 years old to allow plant establishment and inclusion of a range of ages), and 4) shorescape setting as represented by marsh connectivity. For the initial site selection, connectivity was a GIS-derived metric which inversely weighted proximities to natural marsh (positive) and shoreline armoring (e.g,. riprap and bulkhead; negative) such that sites in areas close to marsh and far from armoring would have a high score, and sites in areas close to armoring and far from marsh would have a low score. Candidate sample sites were assigned high, moderate, or low marsh connectivity categories using quantiles in the data distribution (bottom 25% = low, middle 50% = moderate, top 25% = high), a subset of sites was randomly selected from each category, and those that met the above criteria were included in the study; this resulted in near-equal effort in each category (5, 5, and 3 pairs, respectively). Similarly, the sites sampled represented a range of landscape settings (i.e., dominant surrounding land use of agricultural, developed, or natural (i.e., forest, open space) within a 1-km radius) characteristic of Chesapeake Bay (**Table 1**).

**Figure 1.**
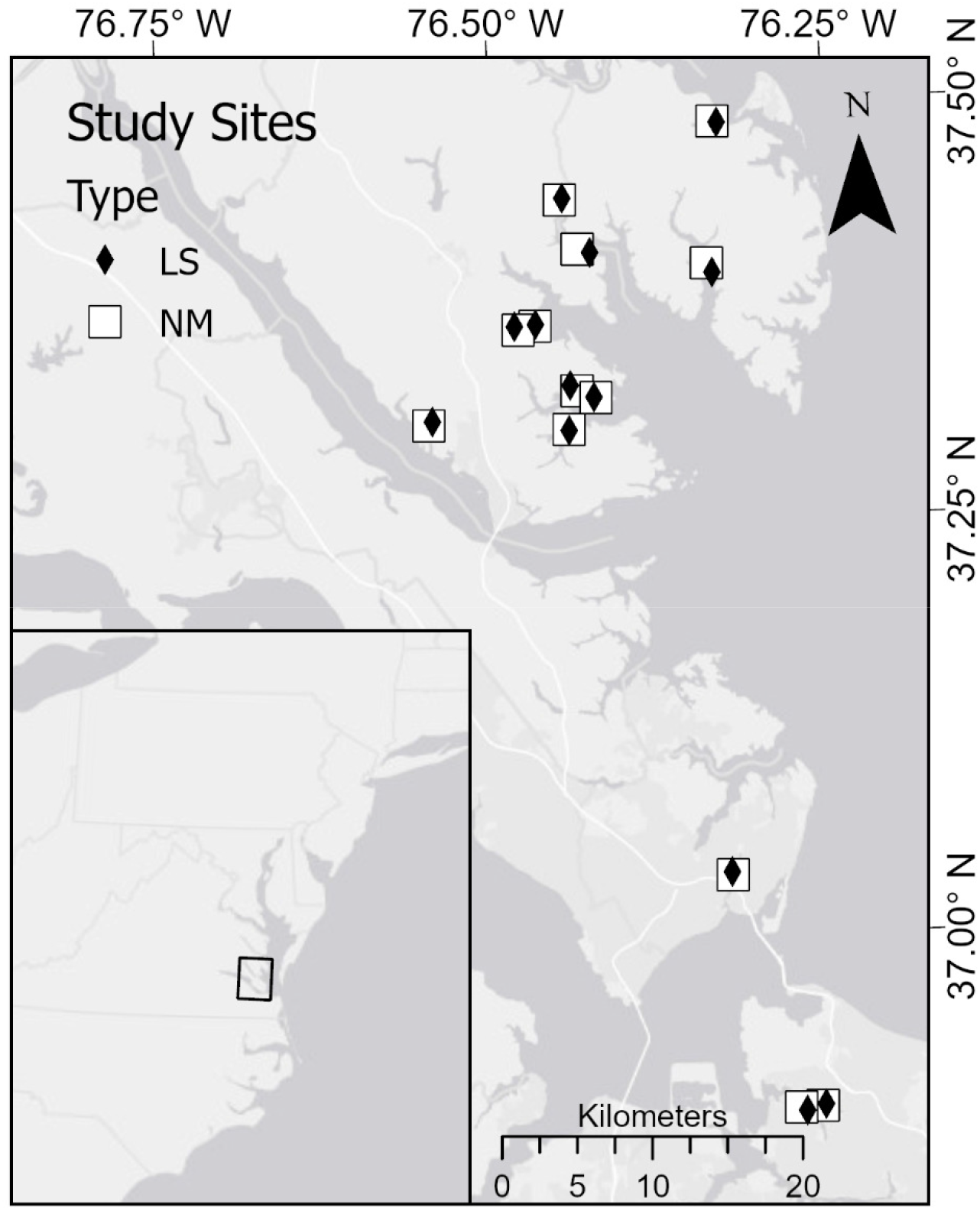
Locations of living shoreline (created fringing marshes) and paired natural fringing marsh study sites in the southern Chesapeake Bay.

**Table 1.** Location, age, inundation period, connectivity, land use characteristics surrounding each living shoreline and natural marsh. Developed, agriculture, managed and natural categories refer to the number of square meters of each land cover type within a 1-km radius of the site; land cover was then classified into 3 major groups: developed (>60% developed and managed lands), natural (>75% natural lands-forest/tree/scrub-shrub), or mixed use (mix of pasture, cropland, developed, managed and natural lands). Shoreline armor refers to the percent armoring within a 1-km radius of the site. Habitat connectivity is represented as the average distance (along the shoreline) to nearest marsh within a 1-km shorescape.

| Pair | Type | Age<br>y | Land<br>cover | Developed<br>(m<sup>2</sup>) | Agriculture<br>(m<sup>2</sup>) | Managed<br>(m<sup>2</sup>) | Natural<br>(m<sup>2</sup>) | Shl<br>armor<br>(%) | Inundation<br>(hrs/mo) | Distance<br>to<br>nearest<br>marsh<br>(m) | Young<br>blue<br>crab<br>density<br>(#/m<sup>2</sup>) |
| --- | --- | --- | --- | --- | --- | --- | --- | --- | --- | --- | --- |
| 1 | LS | 7 | Mix | 94,663 | 870,997 | 479,156 | 1,696,100 | 3% | 146.7 | 6.8 | 0.32 |
| 1 | NM |  | Mix | 69,219 | 809,274 | 462,713 | 1,798,191 | 4% | 285.3 | 5.2 | 0.00 |
| 2 | LS | 4 | Mix | 88,227 | 522,774 | 429,525 | 2,101,070 | 11% | 165.7 | 6.7 | 0.43 |
| 2 | NM |  | Mix | 50,456 | 434,479 | 194,210 | 2,462,449 | 7% | 111.1 | 5.5 | 0.38 |
| 3 | LS | 2 | Nat | 40,143 | 27,130 | 88,183 | 2,986,140 | 0% | 265.1 | 3.6 | 0.86 |
| 3 | NM |  | Nat | 39,548 | 28,347 | 96,151 | 2,977,545 | 0% | 259.4 | 3.5 | 0.14 |
| 4 | LS | 7 | Dev | 1,063,940 | 0 | 872,750 | 1,204,900 | 44% | 141.4 | 11.2 | 0.17 |
| 4 | NM |  | Dev | 1,069,307 | 0 | 846,336 | 1,225,950 | 51% | 97.9 | 14.2 | 0.10 |
| 5 | LS | 10 | Dev | 1,349,630 | 0 | 864,814 | 927,144 | 62% | 93.6 | 28.7 | 0.13 |
| 5 | NM |  | Dev | 1,339,773 | 0 | 957,234 | 844,586 | 62% | 84.1 | 33.9 | 0.58 |
| 6 | LS | 9 | Mix | 72,256 | 380,434 | 175,902 | 2,513,000 | 12% | 142.1 | 5.4 | 1.67 |
| 6 | NM |  | Mix | 83,859 | 506,761 | 202,020 | 2,348,953 | 15% | 209.0 | 5.8 | 1.47 |
| 7 | LS | 12 | Mix | 85,851 | 603,660 | 279,664 | 2,172,420 | 15% | 152.8 | 8.0 | 0.61 |
| 7 | NM |  | Mix | 67,420 | 478,600 | 372,942 | 2,222,633 | 21% | 159.4 | 6.9 | 2.09 |
| 8 | LS | 7 | Mix | 207,183 | 352,958 | 1,032,858 | 1,542,100 | 20% | 110.9 | 28.4 | 0.06 |
| 8 | NM |  | Mix | 208,974 | 197,565 | 1,169,593 | 1,559,446 | 22% | 198.9 | 25.0 | 0.27 |
| 9 | LS | 3 | Mix | 95,919 | 700,849 | 323,144 | 2,021,680 | 6% | 211.7 | 4.2 | 0.84 |
| 9 | NM |  | Mix | 92,039 | 896,230 | 302,020 | 1,851,303 | 6% | 185.4 | 4.3 | 4.69 |
| 10 | LS | 16 | Nat | 113,161 | 174,387 | 463,910 | 2,930,140 | 12% | 224.7 | 4.0 | 3.16 |
| 10 | NM |  | Nat | 110,535 | 112,833 | 465,484 | 2,452,742 | 11% | 18.1 | 3.8 | 1.35 |
| 11 | LS | 9 | Mix | 90,945 | 411,775 | 353,322 | 2,285,550 | 25% | 343.9 | 8.7 | 1.59 |
| 11 | NM |  | Mix | 90,207 | 430,514 | 367,554 | 2,253,317 | 26% | 268.9 | 8.6 | 0.12 |
| 12 | LS | 6 | Dev | 1,171,480 | 0 | 1,270,174 | 699,941 | 41% | 299.7 | 29.5 | 10.82 |
| 12 | NM |  | Dev | 1,161,934 | 0 | 1,312,171 | 667,491 | 43% | 190.2 | 35.9 | 8.33 |
| 13 | LS | 16 | Mix | 168,060 | 430,037 | 1,053,046 | 1,489,860 | 22% | 289.4 | 14.5 | 2.14 |
| 13 | NM |  | Mix | 143,396 | 186,017 | 945,000 | 1,866,609 | 25% | 228.9 | 13.0 | 4.38 |

Living shoreline and paired natural marsh sites (n = 13 pairs) were sampled once during summers (mid-June-Aug) in 2018 and 2019; paired sites were sampled concurrently to ensure similar environmental conditions. Sampling was completed during the period when blue crabs are settling in primary nurseries in Chesapeake Bay (Pile et al. 1996). All sites were not in close proximity to other potential structural nursery habitats (e.g., persistent seagrass beds) to minimize potential confounding factors. At each site, 2 fyke nets were set at high tide and retrieved at low tide; each net fished for 4 hr ± 40 min (SD). Fyke nets were placed at the sill gaps or ends of the living shoreline sites and randomly along the edge of natural marsh sites. At each paired site, fyke net openings were set at same distance from marsh edge ~1 m, depending on sill location relative to the marsh edge). The fyke nets consisted of a 0.9 × 0.9 × 3.0 m compartmentalized, 3.175-mm-mesh bag with 0.9 × 5.2 m wings that stretched out from the bag (set for a total mouth width of 8m) into the marsh. For each sample, all crabs were counted, measured (cw), and then released. The density of blue crabs was determined on the basis of the area of marsh drained by each fyke set. That sampling area was determined with a Trimble Geo 7x handheld by walking the perimeter of the extent of low marsh (*Spartina alterniflora*) being drained into the net; data were converted to areas in ArcGIS Pro.

In addition, local site and shorescape characteristics that may influence marsh habitat use by juvenile blue crabs were collected to inform the analyses, including inundation duration, and physicochemical measures (salinity, water temperature, dissolved oxygen). The duration and extent of inundation of the marsh was estimated to determine the relative amount of low marsh habitat and feeding opportunities available to young blue crabs. We estimated the distribution of tidal inundation across the marsh surface using the closest NOAA tidal predictions (https://tidesandcurrents.noaa.gov/tide_predictions.html) for each reference marsh and living shoreline pair for a one month period of time (July 2018). We used tidal predictions rather than observations from tidal gauges because none of the study sites were adjacent to tidal gauges. Therefore, the tidal analysis allows us to compare general conditions between marsh/living shoreline pairs, but is not an exact accounting of the experienced water levels over the sampling time period (e.g., differences in predicted tides owing to storms are not included in the analysis). Tidal predictions are only furnished as time and height of high and low tides; therefore, we had to interpolate between the high and low water to create a full tidal curve. A frequency analysis was used to calculate the exceedance period at 0.1m increments for each site. Site elevations were obtained using a digital elevation model (DEM) derived from on-site elevation data collected using a stadia rod and a Trimble Geo 7X handheld GPS. The elevations were transformed into hours of inundation using the tidal analysis data and averaged across the low marsh surface for each site. Connectivity as used in the statistical analysis was assessed as the mean distance to natural marsh along a 1-km shorescape such that large values indicate a more isolated site and low values indicate a more connected site. Physicochemical point measures were recorded at the time of sampling for each site using a handheld YSI Exo2.

Mean young blue crab density at a given site was estimated by averaging the 2 fyke net samples for each year. Young blue crab abundance patterns were analyzed using an integrated nested Laplace approximation (INLA; Rue, Martino, & Chopin, 2009) method from the package “INLA” (Martins, Simpson, Lindgren, & Rue, 2013) in R version 4.0.0 (R Core Team 2020). The model structure was specified as:

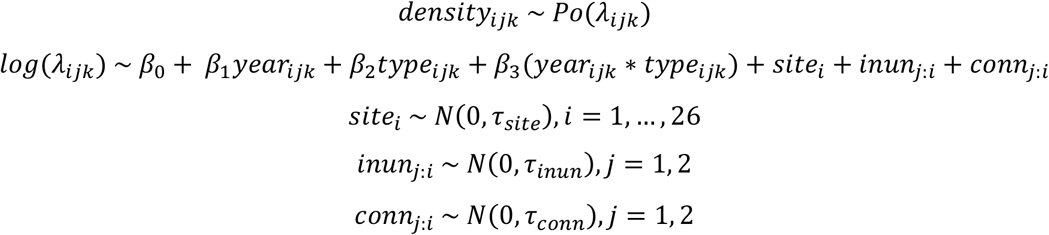

Density of young blue crabs (individuals·m^−2^ of fyke net wing area; multiplied by 10 and integerized) for each site in each year was modeled with a Poisson distribution. Year, type, and their interaction were included as fixed effects, while site, and inundation duration (*inun*) and connectivity (*conn*) both nested within site (*j:i*) were included as random effects. Both inundation duration and connectivity values were centered and scaled. To assess the relative effects of inundation and connectivity on young blue crab density, we averaged the densities of young blue crabs across all fykes in both years for each site. Then, we fit a simple Poisson regression in INLA using the following model structure:

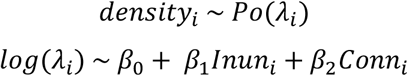

Young blue crab size was similarly analyzed in INLA with the same factors, but with a Gaussian distribution. The model structure was specified as:

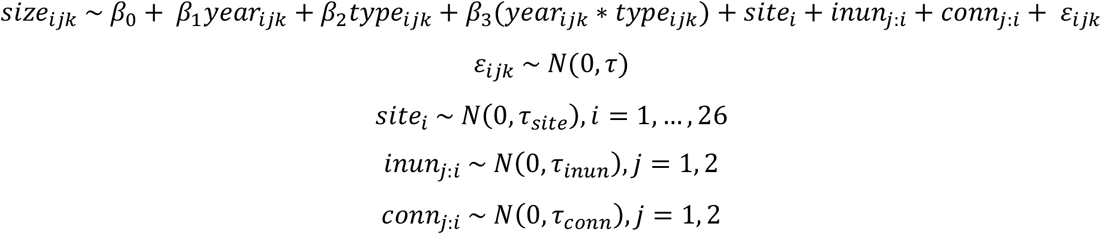

The effects of inundation duration and connectivity on young blue crab size using the following model structure:

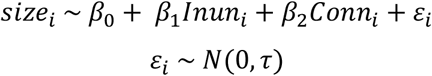

For all INLA mixed-effects models, vague precision priors (gamma distribution: k = 0.001, θ = 0.001) were set for the random effects, and posteriors were computed for each coefficient. The marginal log-likelihood for each model above was compared to the marginal log-likelihood of its corresponding null model to provide a measure of model assessment.

The relationship between the age of the living shoreline and young blue crab abundance was examined with Spearman’s rank correlation ρ.

## RESULTS

A total of 1,704 blue crabs (556 in 2018 and 1148 in 2019) were collected from the 26 sites. Overall, blue crabs were more abundant and smaller on average in 2019 (3.8 cm ± 2.5) than in 2018 (5.3 cm ± 3.3). Of blue crabs collected, a sizable proportion, (24.8% in 2018, 42.8% in 2019) were young juvenile blue crabs (≤ 2.5 cm cw), with peak abundances occurring in August (2018) and July (2019) (**Fig. 2**). Each fyke net sampled an average marsh area of 15.3 ± 2.4 m^2^.

**Figure 2.**
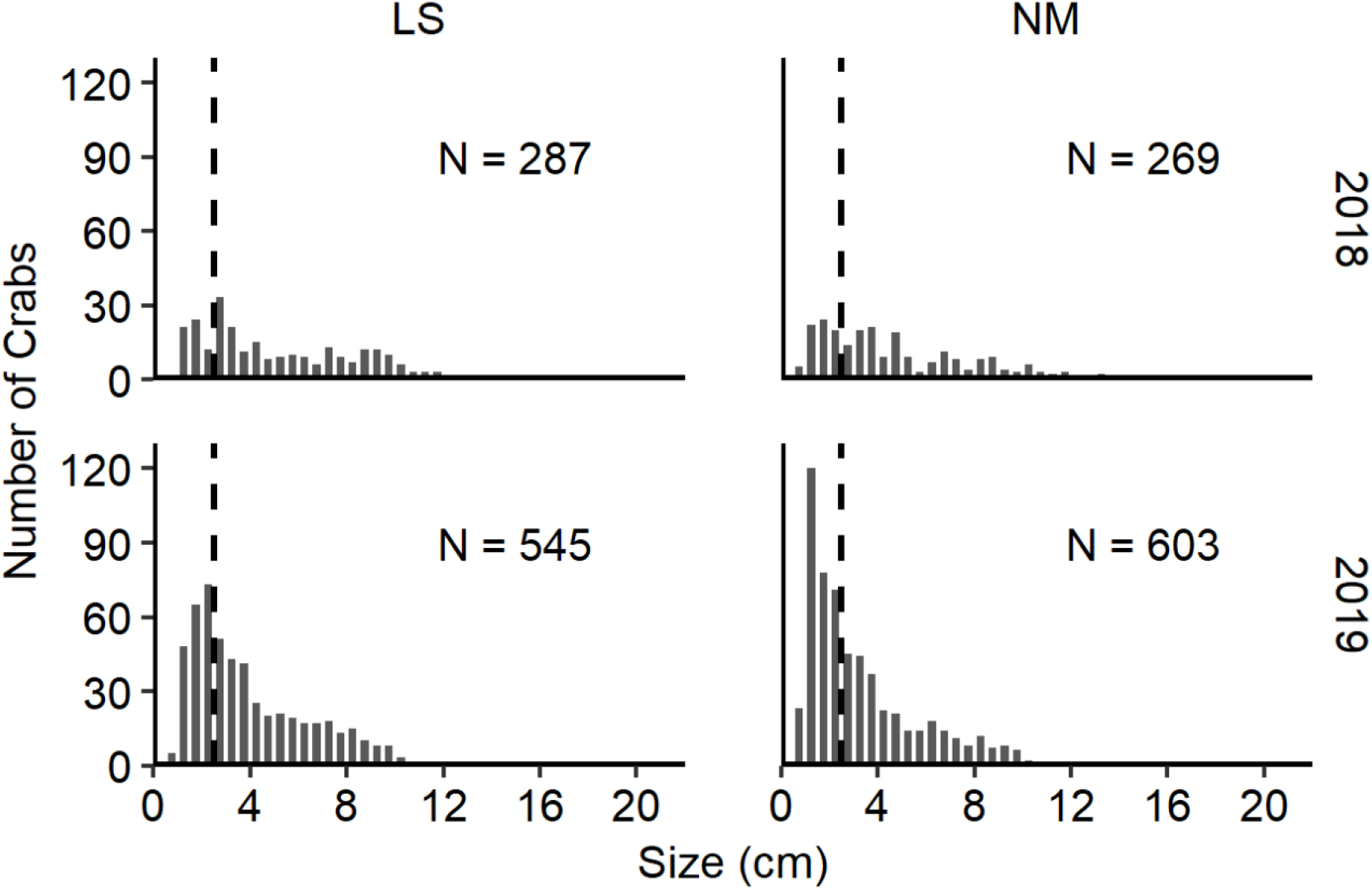
Length-frequency distribution of blue crabs in 2018 and 2019 at (a) living shoreline (LS) created marshes and (b) paired natural fringing marshes (NM) in the southern Chesapeake Bay. Dotted line denotes the approximate size (2.5 cm) when young blue crab juveniles disperse more widely to secondary nursery habitats.

Young blue crabs (≤ 2.5cm) had similar densities (#/m^2^) and sizes in living shoreline and natural fringing marshes (**Table 2**). The only variation in young blue crab densities was related to sample year; in both marsh types densities were greater in 2019 (LS = 0.6 ± 0.2; NM = 0.7 ± 0.2) than 2018 (LS = 0.3 ± 0.2; NM = 0.2 ± 0.1) (INLA Poisson mixed-effects regression, Table 3, **Fig. 3**), although there were no annual differences in mean size (1.8 cm ± 0.4 (2018), 1.7 cm ± 0.5 (2019); **Table 3**). During 2019, annual average freshwater flow into the Chesapeake Bay was the highest on record (since 1937) at 130,750 cubic feet per second (USGS 2020) and this led to a depression in salinity throughout the estuary and at the study sites prior to the sampling period which may have contributed to the annual differences; however, during sampling salinities were similar between years (2018: 15.4 ± 1.9 SD; 2019: 16.1±1.9 SD). Likewise, mean temperature and dissolved oxygen were similar during sampling in 2018 (29.1°C ±1.5 SD; 7.2 mg/L ±1.8 and 2019 (28.3°C ±2.3 SD; 5.6 mg/L ±1.1). Both inundation and connectivity had non-zero effects on young blue crab density, but not on size (**Table 3**). Young blue crab density was higher in marshes and living shorelines that were inundated for longer periods each tidal cycle and was lower in sites located in well-connected shorescapes. Both inundation and connectivity had similar effect sizes.

**Table 2.**
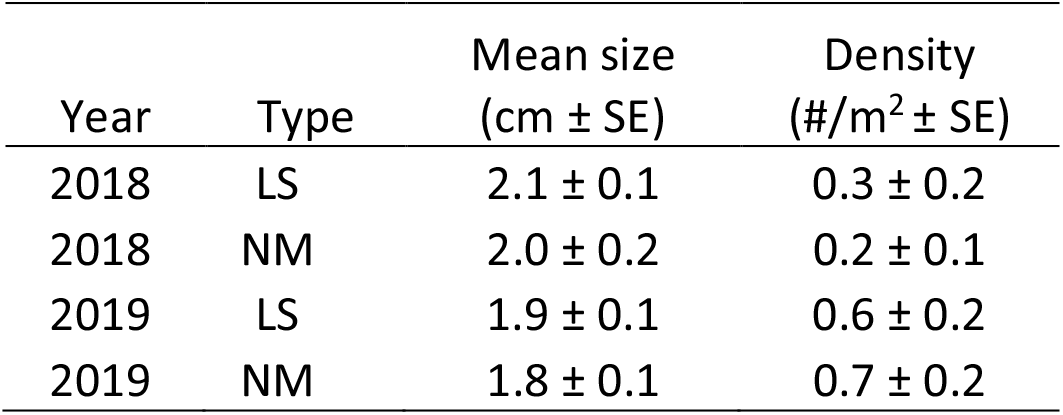
Mean young crab size (cm) and density (individuals·m^−2^) ± standard error (SE) at living shoreline and natural marsh sites for years 2018 and 2019.

| Year | Type | Mean size<br>(cm ± SE) | Density<br>(#/m<sup>2</sup> ± SE) |
| --- | --- | --- | --- |
| 2018 | LS | 2.1 ± 0.1 | 0.3 ± 0.2 |
| 2018 | NM | 2.0 ± 0.2 | 0.2 ± 0.1 |
| 2019 | LS | 1.9 ± 0.1 | 0.6 ± 0.2 |
| 2019 | NM | 1.8 ± 0.1 | 0.7 ± 0.2 |

**Table 3.** Model parameter estimates for the effects of site type and year (Type*Year) models (Eq. 1 & 3) and low-marsh inundation duration and shorescape connectivity (Phys) models (Eq. 2 & 4) on young blue crab density and size. Coefficients indicate the mean response with the 95% credible interval provided in parentheses. Reference levels for the parameters were type = living shoreline and year = 2018. The marginal log-likelihood (MLL) estimates are provided for the given model, with the null model MLL for comparison.

| Model | Intercept | Type | Year | Type*Year | Inundation | Connectivity | MLL | Null |
| --- | --- | --- | --- | --- | --- | --- | --- | --- |
| Density<sub>(Type*Year)</sub> | -0.045<br>(-1.109, 0.930) | -0.326<br>(-1.845, 1.170) | 0.878<br>(0.476, 1.301) | 0.500<br>(-0.105, 1.113) | - | - | -156.87 | -291.70 |
| Size<sub>(Type*Year)</sub> | 2.094<br>(1.770, 2.416) | 0.001<br>(-0.461, 0.463) | -0.153<br>(-0.495, 0.194) | -0.112<br>(-0.587, 0.383) | - | - | -61.10 | -30.46 |
| Density<sub>(Phys)</sub> | 1.910<br>(1.747, 2.066) | - | - | - | 0.541<br>(0.407, 0.677) | 0.550<br>(0.431, 0.668) | -163.22 | -210.75 |
| Size<sub>(Phys)</sub> | 1.960<br>(1.827, 2.092) | - | - | - | -0.036<br>(-0.174, 0.102) | -0.076<br>(-0.211, 0.059) | -28.77 | -17.21 |

**Figure 3.**
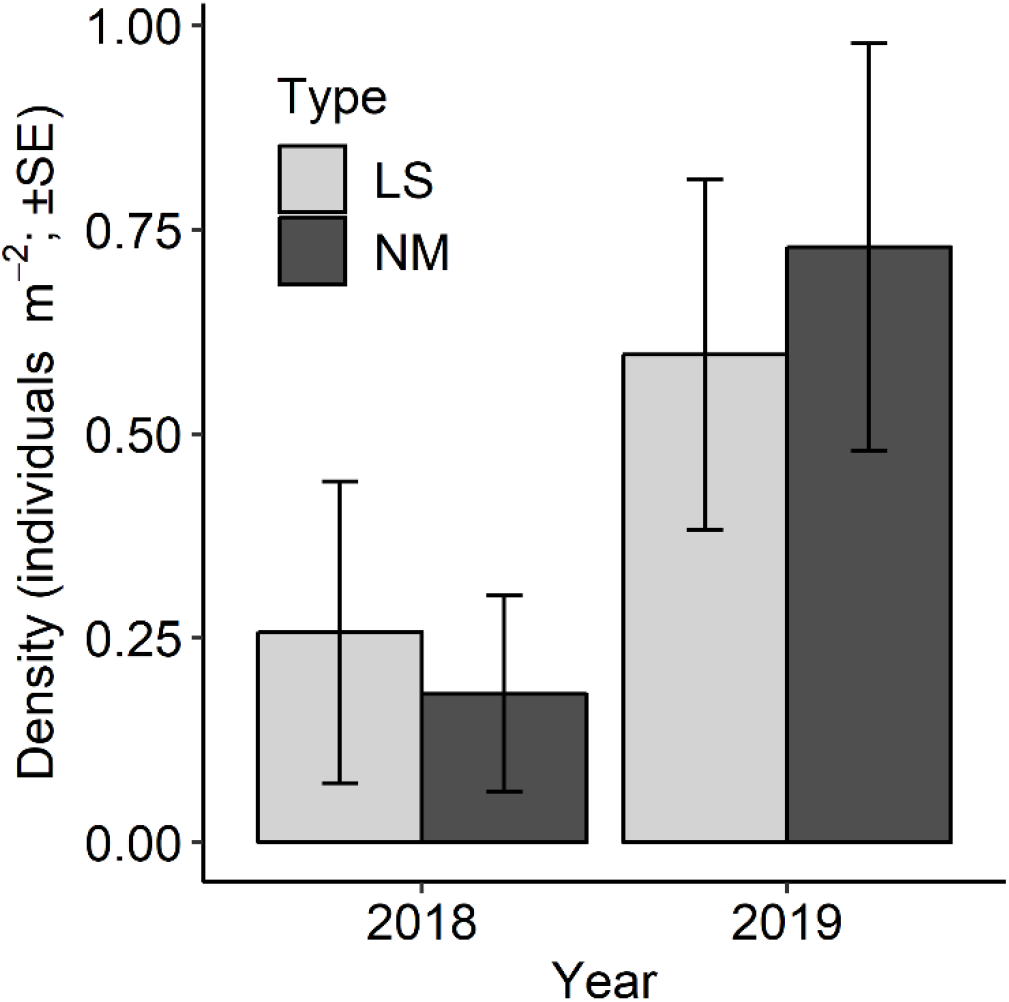
Juvenile blue crabs (≤ 2.5cm) had similar densities (#/m^2^) in living shoreline and natural fringing marshes during each year, with greater densities in 2019 compared to 2018.

There was no relationship between the age of the created marsh and young blue crab density (ρ = 0.22, *p* = 0.48; **Fig. 4**), and similar densities between living shoreline and natural marshes indicated that comparable habitat use occurred within 2 years of marsh creation (the youngest marsh sampled). There was also no evident relationship between the mean size of young blue crabs and sampling date (**Fig. 5**).

**Figure 4.**
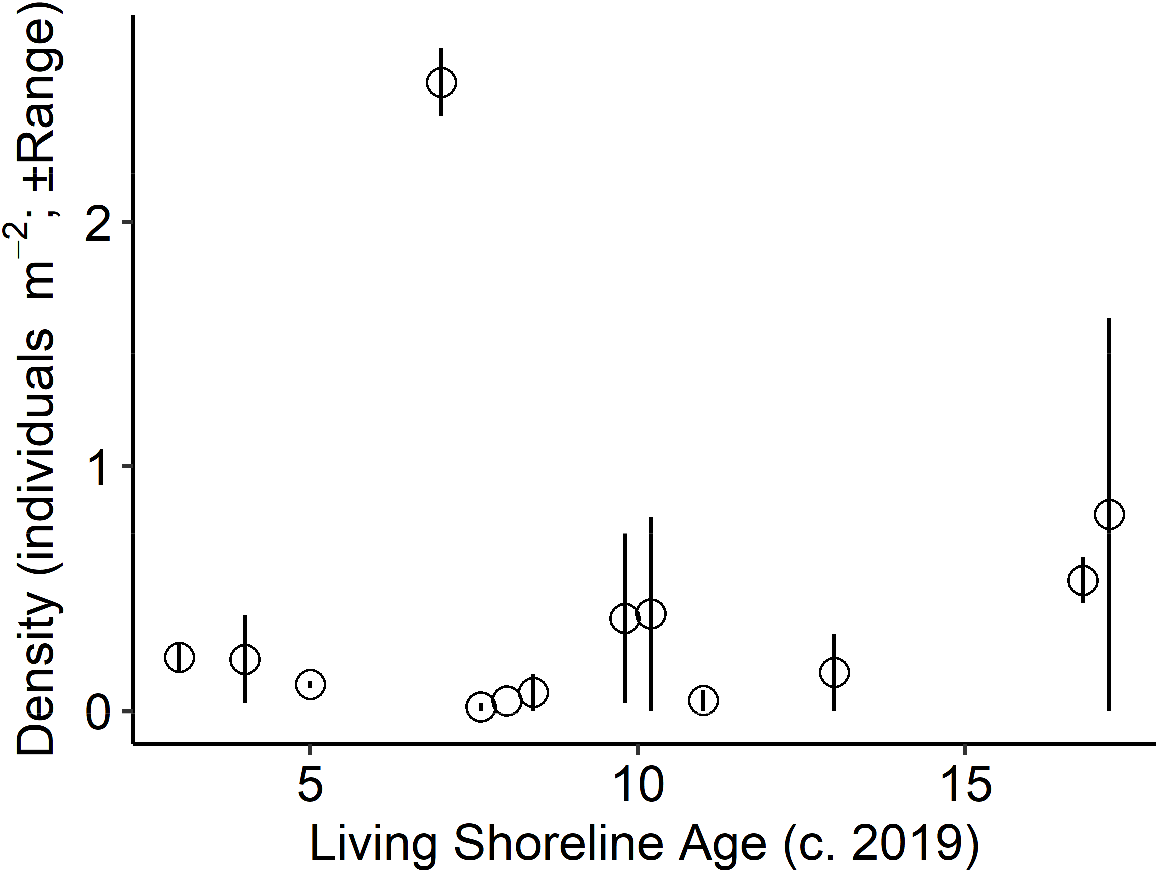
There was no evident relationship between the age of the created marsh and the abundance of young blue crabs, with similar densities of crabs occurring along the chronosequence of marsh ages (2 to 16 years). A small jitter was added along the x-axis for sites with the same age to facilitate visualization (i.e., three sites were 8 Y/O, two sites were 10 Y/O, and 2 sites were 17 Y/O in 2019).

**Figure 5.**
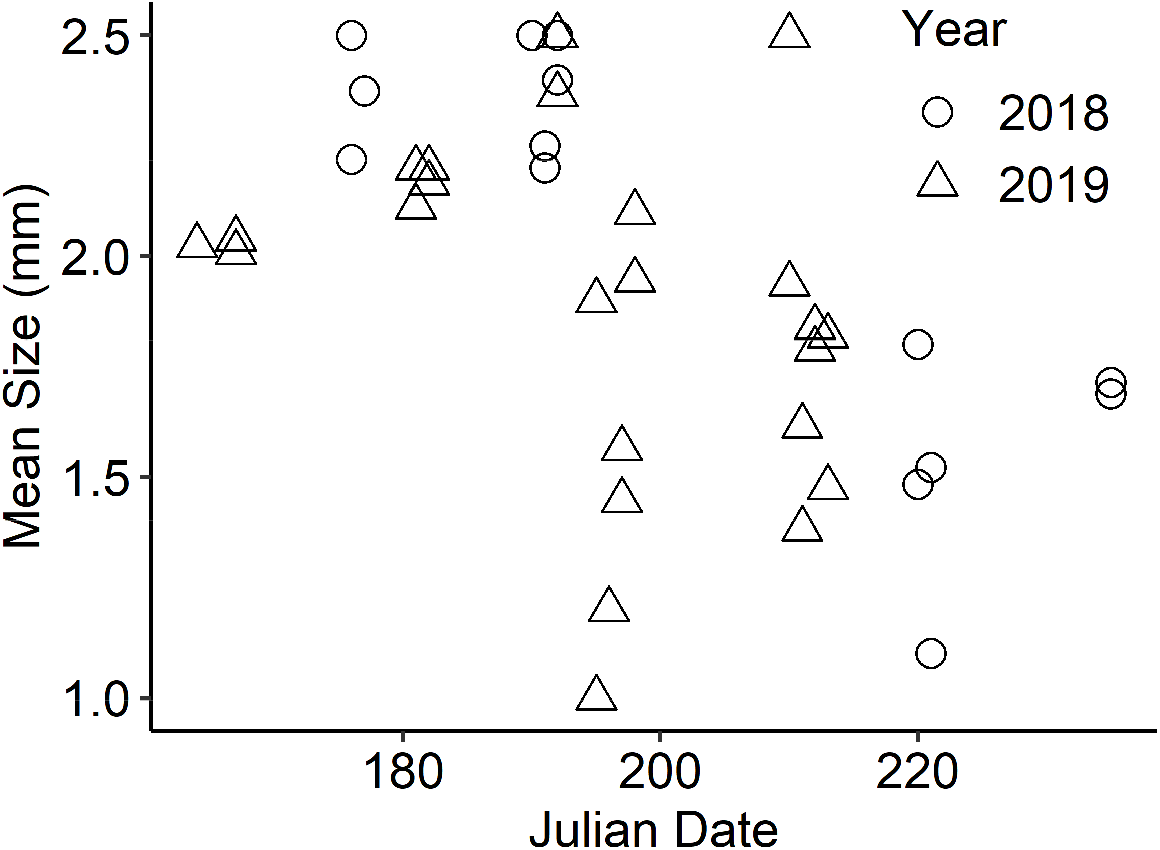
There were no evident seasonal shifts in blue crab size (carapace width in mm) during the sampling periods, suggesting that recruitment was occurring throughout the sampling window (June-August).

## DISCUSSION

Young blue crabs (≤ 2.5cm) were observed in similar abundance and with similar sizes within fringing natural and created marshes suggesting that these marshes are being used as primary nursery habitats. Prior studies have documented living shoreline marsh use by nekton including blue crabs (e.g., Balouskus & Targett, 2016; Gittman et al., 2016), but this study is the first quantification of young blue crab use of living shoreline marshes in Chesapeake Bay. There were annual differences in abundance of blue crabs with higher abundance in 2019; however, these differences were consistent between natural and created marshes. The higher abundance of young blue crabs in 2019 compared to 2018 was consistent with abundance estimates of age 0 crabs from the Chesapeake Bay winter dredge survey which is used to estimate blue crab population health (CBSAC 2019). In the 2018 survey, the abundance of age 0 crabs (< 60mm cw) was below the time series (1990-2019) average of 224 million crabs, and in 2019, the estimate was well above the average at 323 million crabs, which was in the top 27% of time series estimates (CBSAC 2019). We saw no evident seasonal shifts in blue crab size during the sampling periods (**Fig. 5**), suggesting that recruitment was occurring throughout the sampling window (June-August).

The value of marsh edge habitat for fish and invertebrates is well established (Beck et al., 2001; Minello et al., 2003), and preferential habitat use by fish and crustaceans appears to be similar along the edges of narrow fringing and extensive marshes (Minello, Zimmerman, & Medina, 1994; Peterson & Turner, 1994; Micheli & Peterson, 1999; Currin, Delano, & Valdes-Weaver, 2008). In addition, juvenile blue crabs and a variety of commercially or ecologically important finfish exhibit a strong preference (higher abundance and/or biomass) for fringe marsh habitat over altered shoreline (e.g., Peterson, Comyns, Hendon, Bond, & Duff, 2000; King et al., 2005; Bilkovic & Roggero, 2008; Balouskus & Targett, 2016). Less understood is the relative importance of salt marshes as young crab nursery habitat, which may vary among systems depending on the composition of marine coastal habitats and environmental conditions (Johnson & Eggleston, 2010; Shakeri, Darnell, Carruthers, & Darnell, 2020).

Examining the relative densities of juvenile blue crabs in different potential nurseries is one way to evaluate the relative importance of each habitat. Young blue crab density within seagrass beds in Chesapeake Bay decreases with increasing instar stage (third to the seventh instar), possibly because crabs are migrating from seagrass habitat into other nursery habitats, particularly salt marshes (Pile, Lipcius, Van Montfrans, & Orth, 1996). The marsh edge densities from this study were generally lower than those observed in seagrass beds for 7^th^ to 9^th^ instars (^~^10.7 to 16.1 mm; ^~^0 to 5 individuals/m^2^, interpreted from Figure 4 in Pile et al. 1996). However, this is an imperfect comparison as our study likely underestimates crab density within vegetated marsh edges, because the marsh area sampled encompasses habitat that is likely sub-optimal for blue crabs (i.e., inner marsh surface). Also, blue crabs around the 7^th^ instar (~10 mm) are at the limits of detection for the sampling gear (3.175-mm-mesh fyke net); whereas, Pile et al. (1996) sampling gear targeted 1^st^ through 9^th^ instars (2.2 to 16.1 mm). Additional research is needed to quantify the comparative occupancy on earlier instar stages within marshes.

The amount and spatial extent of marshes may exceed other marine habitats that serve as blue crab nurseries (e.g., *Zostera*/*Ruppia* seagrass meadows) which have more restricted distributions (e.g., narrow salinity range, temperature limitation) or high annual variability. This may be particularly evident at the limit of species ranges. In Chesapeake Bay, for example, eelgrass *Zostera marina* is at the southernmost extent of its range and highly susceptible to prolonged high summer temperatures (Orth et al., 2017). Further, the total tidal emergent salt and brackish marsh area in Chesapeake Bay is 1438 km^2^ (NWI Version 2, downloaded 2019; CCRM, 2017), whereas submerged aquatic vegetation area (excluding tidal fresh) on average (2009–2018) is 304 km^2^ (VIMS SAV Program, http://web.vims.edu/bio/sav/ZoneDensityTable.htm). Thus, the value of marshes as primary nursery habitats for blue crab likely varies temporally and spatially; in years and areas with poor seagrass production because of physical conditions, the primacy of marshes as nursery habitat may occur. Alternately, in areas with expansive persistent seagrasses, marshes may be less important to young recruits. Fringing marshes are a geomorphic subcategory of tidal marshes that may be particularly important as blue crab nurseries (Epifanio, 2019) and these narrow bands of vegetation (marsh edge) can make up large proportions of an estuarine shoreline. In Chesapeake Bay, these fringing marshes occupy 15% (~ 2,800 km) of the total 19,000 km of Chesapeake Bay tidal shoreline (CCRM, 2019). The creation or restoration of fringing marshes as living shorelines could help enhance this nursery habitat for blue crabs. Ultimately, a suite of nursery habitats may be essential for long-term successful recruitment in environmentally variable systems.

Fringing marshes may serve to connect other estuarine habitats and habitat complexes (Able, Vivian, Petruzzelli, & Hagan, 2012; Davis, Johnston, Baker, & Sheaves, 2012) that would otherwise be isolated by coastal development; therefore, we posited that the level of marsh connectivity may influence the distribution and abundance of young crabs. In Coastal Virginia, marsh connectivity did have an impact on juvenile blue crab density, though not in the direction that we initially expected. The negative association between young blue crab density and proximity of other marshes within the shorescape may point to an increased encounter rate in poorly connected marshes. When refugia are limited within a system, individuals may pack more densely into the available habitat, thereby increasing observed densities (Bohnsack, 1989). Therefore, we do not conclude that poorly-connected marshes are better for young blue crabs than well-connected marshes, but instead suggest that the higher observed densities in more isolated marshes adds further support to the importance of fringing marshes as nursery habitat for young blue crabs in Chesapeake Bay. This also suggests that crabs are not randomly distributed, but are using cues to settle in these isolated marshes. Of similar importance to juvenile crab abundance was the geomorphology of the fringing marshes. Those marshes that were submerged longer had higher juvenile blue crab densities, likely reflecting the relative availability of refugia and feeding opportunities for the young crabs. Likewise, those living shoreline marshes built higher in the tidal envelope provided less habitat for juvenile crabs, an important consideration when designing living shorelines to enhance habitat provision services.

Managing for multiple connected nursery habitats across seascapes may enhance the resilience of blue crab populations to climate and human pressures (e.g., Boström, Pittman, Simenstad, & Kneib, 2011; Pittman, Kneib, & Simenstad, 2011; Nagelkerken, Sheaves, Baker, & Connolly, 2015). Marshes will be able to withstand prolonged periods of high temperatures that lead to seagrass dieback in the Chesapeake Bay, and may increasingly serve as critical nurseries within a suite of habitats that support the growth and survival of blue crab under changing climate. Climate adaptation strategies that include restoration of marsh shorelines (living shorelines) could not only reduce erosion, flooding, and enhance resilience, but also provide essential habitat for blue crabs. This study demonstrated that within two years of restoration, living shorelines are being used by young blue crabs at similar levels as natural fringing marshes. Much of the eroding Chesapeake Bay shorelines are suitable for the construction of a living shoreline (Berman, Mason, Nunez, Tombleson, 2018; Bilkovic, Mitchell, Havens, Hershner, 2019); yet the implementation of these projects is still relatively low compared to shoreline armoring (bulkhead; riprap revetments). However, there has been growing acceptance and implementation of these projects in Chesapeake Bay and along the Atlantic and Gulf Coasts, coinciding with the blue crab range. Furthermore, living shoreline marshes that enhance sediment capture rates (Currin et al., 2008) and dissipate wave energy protecting the sediment below the root depth, may be able to persist under sea level rise conditions longer than natural marshes (Mitchell & Bilkovic, 2019). This study documents another significant co-benefit of living shorelines – nursery habitat of an important commercial species – that could be used in further justification/valuation of restoration activities.

## Authors’ contributions

Bilkovic conceived of and designed the study, completed field collection, and conducted data interpretation. Chambers, Stanhope, and Angstadt completed most of the field collection; Chambers contributed to data interpretation; Isdell conducted statistical analyses, developed marsh connectivity and inundation analytical approaches, and contributed to data interpretation; Havens assisted with field collection and data interpretation. *All authors contributed critically to the drafts and gave final approval for publication*.

## Acknowledgments

Thanks to all the homeowners who provided access to their marshes used in this study, and CCRM scientists for their critical assistance with sample collection. Molly Mitchell provided conducted tidal predication analyses for the marsh inundation estimates. This material is based upon work supported by the National Science Foundation under Grant Number 1600131, Coastal SEES Collaborative Research Program: Sustainability in Chesapeake Bay shorescapes. Any opinions, findings, and conclusions or recommendations expressed in this material are those of the authors and do not necessarily reflect the views of the National Science Foundation. This paper is Contribution No.xxxx of the Virginia Institute of Marine Science, William & Mary.

## Data Accessibility Statement

Data will be made available from the W&M ScholarWorks Repository, https://scholarworks.wm.edu/.

